# Neural correlates of crowding in macaque area V4

**DOI:** 10.1101/2023.10.16.562617

**Authors:** Taekjun Kim, Anitha Pasupathy

**Affiliations:** Department of Biological Structure, University of Washington, Seattle, WA 98195; Washington National Primate Research Center, University of Washington, Seattle, WA 98195

## Abstract

Visual crowding refers to the phenomenon where a target object that is easily identifiable in isolation becomes difficult to recognize when surrounded by other stimuli (distractors). Extensive psychophysical studies support two alternative possibilities for the underlying mechanisms. One hypothesis suggests that crowding results from the loss of visual information due to pooled encoding of multiple nearby stimuli in the mid-level processing stages along the ventral visual pathway. Alternatively, crowding may arise from limited resolution in decoding object information during recognition and the encoded information may remain inaccessible unless it is salient. To rigorously test these alternatives, we studied the responses of single neurons in macaque area V4, an intermediate stage of the ventral, object-processing pathway, to parametrically designed crowded displays and their texture-statistics matched metameric counterparts. Our investigations reveal striking parallels between how crowding parameters, e.g., number, distance, and position of distractors, influence human psychophysical performance and V4 shape selectivity. Importantly, we found that enhancing the salience of a target stimulus could reverse crowding effects even in highly cluttered scenes and such reversals could be protracted reflecting a dynamical process. Overall, we conclude that a pooled encoding of nearby stimuli cannot explain the observed responses and we propose an alternative model where V4 neurons preferentially encode salient stimuli in crowded displays.

**Significance Statement:** Psychophysicists have long studied the phenomena of visual crowding, but the underlying neural mechanisms are unknown. Our results reveal striking correlations between the responses of neurons in mid-level visual cortical area V4 and psychophysical demonstrations, revealing that crowding is influenced not only by the number and spatial arrangement of distractors but also by the similarity of features between target and distractors, as well as among the distractors themselves. Overall, our studies provide strong evidence that the visual system uses strategies to preferentially encode salient features in a visual scene presumably to process visual information efficiently. When multiple nearby stimuli are equally salient, the phenomenon of crowding ensues.

## Introduction

In cluttered natural visual environments, object recognition capacity can be severely limited. This is well-demonstrated by find-the-difference puzzles and inattention blindness displays where even large objects can be missed (1–3). Human psychophysical studies have explored this issue in detail for decades, and have demonstrated that diminished object recognition in clutter, referred to as “crowding”, has several defining characteristics (4–7). We now know that the discriminability of an object (i.e., target) depends on the distance between that object and surrounding distractors: nearby distractors more severely limit discriminability. We also know that crowding worsens with distance from the fovea: target–distractor distance over which crowding operates increases with eccentricity. Third, at a fixed target-distractor distance, the crowding zone is not circular but elongated toward the point of fixation, such that radially positioned distractors induce a stronger crowding effect than tangentially positioned ones (8–10).

Crowding may represent the loss of visual information, a by-product of how visual stimuli are encoded along the successive stages of the ventral visual pathway (11–13). One concrete hypothesis posits that single neurons in mid-level stages of the ventral stream (e.g., V2, V4) encode the aggregated statistics of the image region within their receptive fields (the texture-pooling model) resulting in crowding (14–16). Alternatively, the encoding process may not be lossy; instead, crowding may result from limited resolution of object decoding during recognition, i.e., encoded information may remain inaccessible unless salient (17–20). However, neurophysiological studies have seldom queried the responses of neurons to crowded displays and the neuronal mechanisms of crowding remain unknown.

In this study, we targeted macaque area V4, an intermediate stage in the ventral visual pathway known to be critical for object shape processing and recognition (21). We used parametrically designed arrays of shape stimuli to investigate how shape selective responses of V4 neurons, in two awake, fixating macaque monkeys, were influenced by the manipulation of distance, number, and position of clutter stimuli. Across the V4 population, we evaluated whether the most effective distractor position varies based on the receptive field location, as predicted by anisotropy of crowding zones, providing a neurophysiological correlate to human psychophysical observations. We also sought to assess the appropriateness of the texture pooling model by studying neuronal responses to texture-statistics-matched metameric stimuli and crowded displays where we titrated target saliency. To gain insights into the temporal dynamics of representational processes that impact visual crowding, we conducted single trial population decoding analyses as a function of time. Lastly, based on our findings, we propose a novel model that selectively processes salient features in a cluttered visual scene to efficiently manage visual information.

## Results

We studied the responses of 147 V4 neurons in two awake, fixating macaque monkeys (66 in Monkey 1, 81 in Monkey 2) to examine how their responses and selectivity to an isolated shape are modified by the presence of surrounding clutter stimuli. We used a large set of systematically designed stimulus arrays (Figure 1) where we varied the number, distance and position of the distractor stimuli, and the salience of target stimuli, to probe how V4 responses correlate with the perceptual characteristics of crowding.

**Figure 1.**
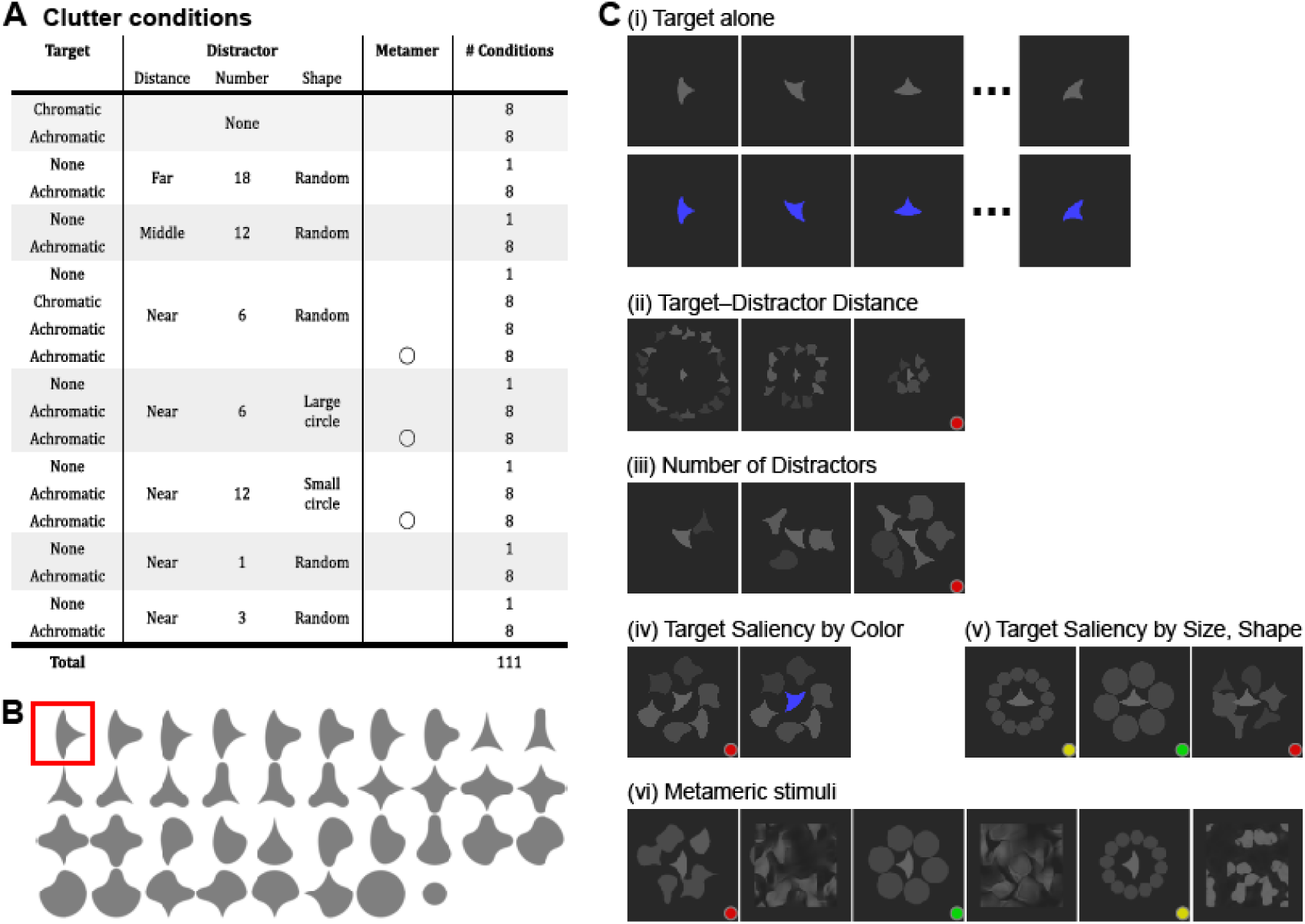
Visual stimulus design. **A.** Tabulation of clutter conditions. For each target– distractor configuration (rows), responses to eight target rotations (# Conditions = 8) were evaluated, except when distractors were presented without a target (Target = None). An open circle under *Metamer* column indicates that metameric stimuli were also presented (see Methods). **B.** Shape stimulus set. Target (red box) and distractor shapes were chosen from a set of 2D shapes (a subset of shapes created by (22)). Target stimulus was at the RF center, scaled to half of the estimated RF diameter. **C.** The target stimulus was presented either (i) alone or in combination with various distractor arrangements which varied in terms of (ii) distance from central target, (iii) number, (iv) saliency defined by target color, (v) shape, shape + size of distractors. In all conditions, the target shape was shown at eight rotations in 45° increments (as in (i)). Targets were achromatic or chromatic when presented alone. In all clutter conditions targets were achromatic except when target saliency was titrated by color (iv). Distractors were always achromatic. The target size was the same in all conditions, but it is scaled down in (ii) for illustration purposes. Distractors were the same size as target except when titrating saliency by size (yellow dot). Metameric stimulus pairs with matched texture statistics (in (vi): panels 1-2, 3-4 and 5-6) were included to test the texture pooling model (see Methods). Colored dots identify identical stimulus conditions repeated in the figure for illustration purposes.

### Effects of target-distractor distance on target shape selectivity

A fundamental psychophysical observation is that crowding effects dissipate with increasing distance between target and distractors (4, 5, 7). To determine if V4 responses exhibit a similar trend, we examined how shape selectivity and response strength are modulated by distractors at three different distances from the target stimulus centered within the RF. Results from an example neuron are shown in Figure 2A-C. When the target stimulus was presented in isolation, this neuron showed diverse responses across stimulus orientation (Figure 2A, red). When the distractors were far from the target (Figure 2A, 2^nd^ row), responses were weaker than for target alone (distractor modulation index = −27%), especially for the preferred target orientations (compare columns 1-4 for rows 1 and 2). Despite this, the responses showed considerable variability across target orientations (one-way ANOVA, p < 0.01) and the overall tuning curve was similar to that when the target was in isolation (compare red and light gray curves, Figure 2B), implying that shape selectivity was well maintained (correlation coefficient (*r*) = 0.97, p < 0.01) despite suppressive modulation by surrounding distractor stimuli. Results were similar when the distractors occupied an intermediate distance (see Figure 2A, 3^rd^ row; Figure 2B, dark gray). However, when the distractors were close to the target, the variation in responses across target orientation was weak (compare black vs. other lines in Figure 2B), and as expected, there was weak correlation between the tuning curves for target alone and target + near distractors (*r* = 0.28, p = 0.49).

**Figure 2.**
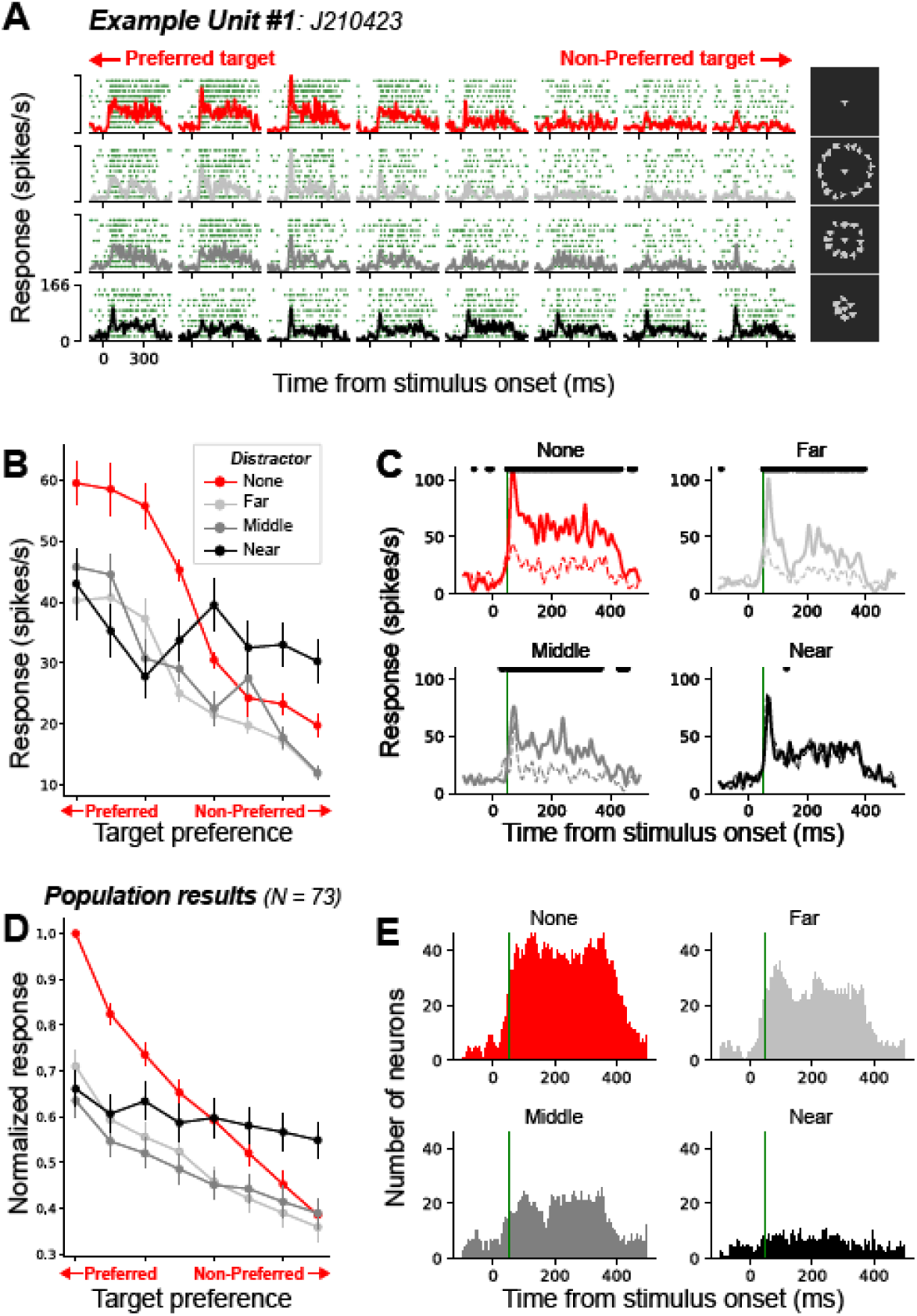
Effect of target-distractor distance on shape tuning. **A-C. Example neuron. A.** Raster plots and PSTHs for responses to: (rows) target alone, target + far, middle, and near distractor conditions, respectively. Columns show responses to different target orientations, rank-ordered by responses to target-alone. Stimulus panels are shown here at a higher contrast to aid visibility (see Figure 1C for veridical illustration). **B.** Tuning curves based on average responses (0-400 ms) for the four target + distractor conditions. Error bars indicate the standard error of the mean. **C.** Average PSTHs for the preferred (top 4) and non-preferred (bottom 4) targets for each distractor condition are shown in solid and dotted lines, respectively. Black asterisks indicate time points with significant difference between solid and dotted curves (Mann–Whitney U test in a 30 ms sliding window, p < 0.05). **D-E.** Population results. **D.** Average normalized tuning curves across the sub-population of neurons with significant shape selectivity in the target alone condition (73/147). Error bars indicate the standard error of the mean. **E**. The number of significantly modulated neurons as a function of time for each distractor condition. Green vertical lines at 50 ms after the stimulus onset are included to facilitate comparison across panels.

Figure 2C illustrates the time course of target shape selectivity as a function of distractor distance. For far and intermediate distractor distance, statistically significant difference between preferred (solid lines) and non-preferred (dotted lines) targets emerged early (soon after visual response onset) but this was not the case for near distractors.

We observed similar results across our sub-population of 73 shape selective neurons. Population average tuning curves for target + distractor conditions were shallower (Figure 2D) and correlation with target alone decreased (median correlation coefficient (r): 0.82 → 0.69 → 0.44) as target–distractor distance decreased. Differences in all pairwise comparisons between three distributions were statistically significant (Wilcoxon signed-rank test, p < 0.01). The magnitude of suppressive modulation by surrounding distractors was also strongest in the near distractor condition (46% decrease) and weaker in the far distractor condition (32% decrease) (Wilcoxon signed-rank test, p < 0.01). However, in each of the three distance conditions, the degree of visual crowding (i.e., impaired shape selectivity, quantified by Fisher’s z-transformed correlation in responses between target alone and target + distractors) was not correlated with the amount of surround modulation (i.e., the absolute value of modulation index) (Far: r = −0.18, p = 0.12; Middle: r = −0.13, p = 0.27; Near: r = 0.16, p = 0.18; data not shown), possibly suggesting that suppression and crowding are separate mechanisms. The time course of emergence of target shape selectivity across the population was also very similar for target alone and target + far/middle distractor conditions (Figure 2E): for all except the near distractor case, the number of significantly modulated neurons showed a sharp rise ∼50 ms after the stimulus onset (green vertical lines).

### Effects of distractor number on target shape selectivity

Psychophysical studies have also documented that the effects of crowding increase with the number of distractors. To assess the influence of the number of distractors on V4 shape selectivity, we compared responses in the ‘target alone’ condition with those from target + one, three, and six distractor conditions. In this case, all distractors were placed at the same distance from the target (i.e., 0.5 × RF diameter, Figure 3A). Results from an example neuron (#2, Figure 3) shows that target shape preference was consistent between the ‘target alone’ and ‘target + one distractor’ conditions (Figure 3A-B, correlation coefficient (r) = 0.90, p < 0.01). However, as the number of distractors increased, shape dependent response variation gradually decreased, and shape selectivity could no longer be observed in the ‘target + six distractor’ condition (one-way ANOVA, p = 0.78). This was consistent with the reduced separation between the average PSTHs for preferred (solid line) and non-preferred (dotted line) targets with increasing distractor number (Figure 3C).

**Figure 3.**
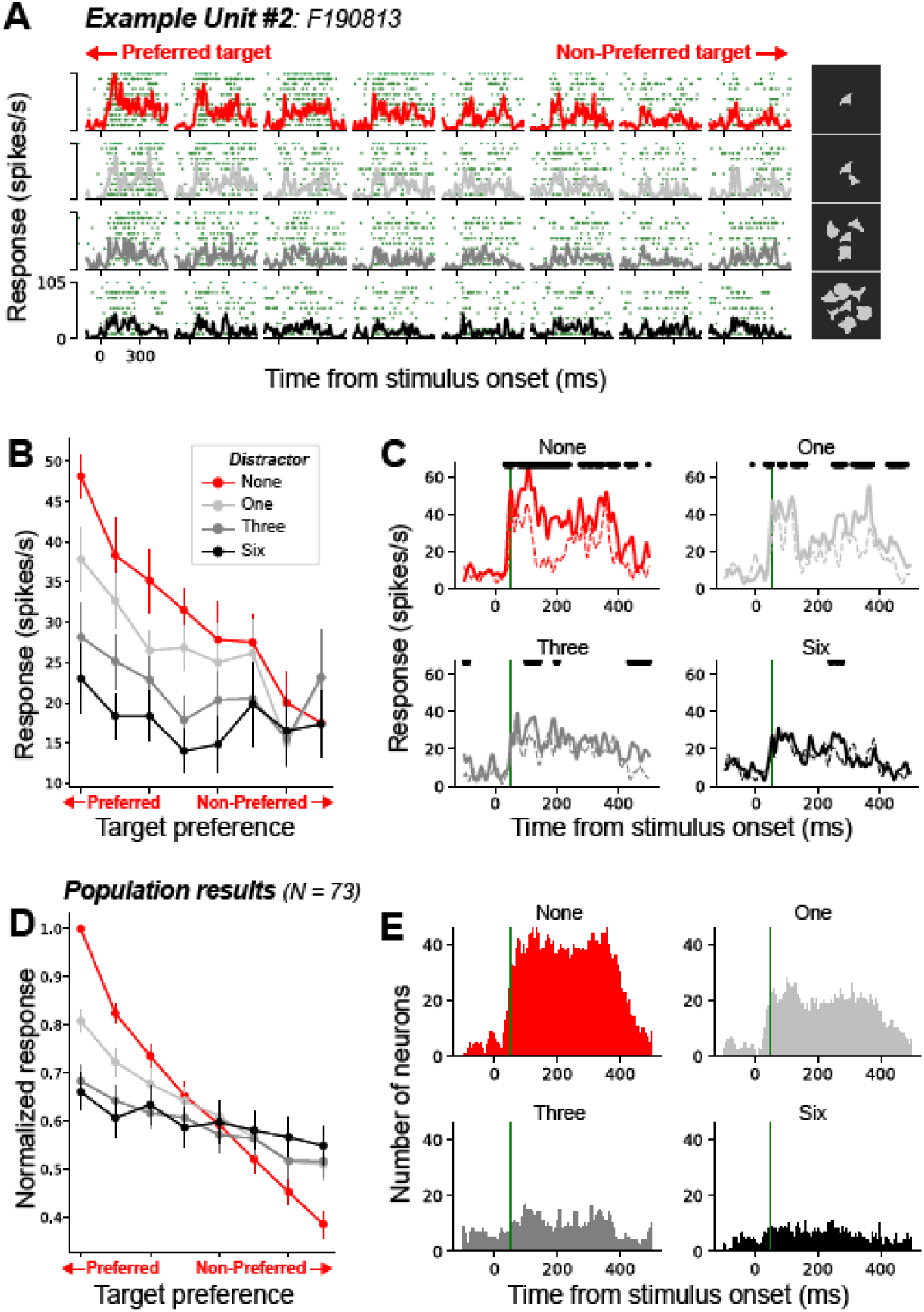
Effect of distractor number on responses and selectivity. **A-C.** Example neuron responses. **A.** Raster plots with PSTHs for responses to: (rows) target alone, target + 1, 3 or 6 distractors, respectively. Columns show responses to different target orientations. **B.** Target shape selectivity curves of the example unit for the four different conditions. **C.** Average PSTHs for the preferred (top 4) and non-preferred (bottom 4) targets. **D-E.** Population results. **D.** Average normalized tuning curves for target shape selectivity in the presence and absence of distractors. **E.** The number of significantly modulated neurons as a function of time for each distractor condition. All conventions are as in Figure 2.

Population results confirm that more numerous distractors generally induce stronger suppression of preferred responses and a decline in shape selectivity (Figure 3D-E). As the number of distractors increased (1 → 3 → 6), the median correlation coefficient with target alone decreased (median (r): 0.79 → 0.54 → 0.44). The median values of suppressive modulation in one, three, six distractor conditions gradually increased from −37% to −42% to −46%, respectively. A noteworthy feature is that the strength of suppression from distractors did not increase linearly: three or six-times as many distractors did not result in three- or six-times stronger surround suppression.

### Anisotropy of visual crowding in area V4

In addition to distance and number effects of distractors, psychophysical experiments have reported significant inward-outward anisotropy in visual crowding (23, 4, 24). Along a radial axis, connecting target position and the fixation point, studies have found a greater interference from distractors positioned beyond the target than between the target and the fixation point (see Figure 4A). To examine whether V4 provides a neural correlate for this psychophysical finding, we compared the strength of visual crowding effect across various near-surround positions.

**Figure 4.**
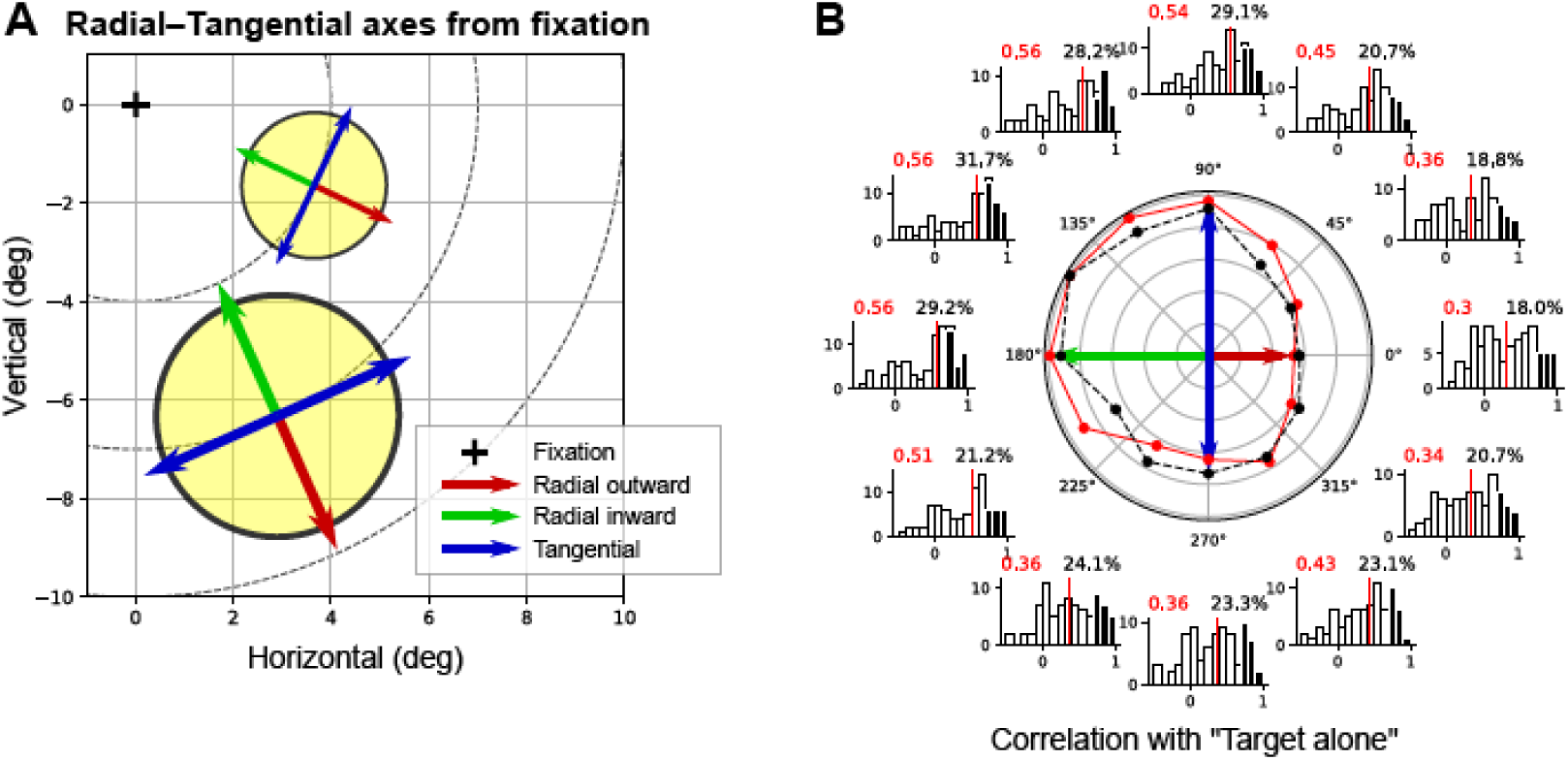
Effects of distractor position on visual crowding: anisotropic crowding zone. **A**. For each neuron, the positions of the distractors were realigned based on the radial and tangential axes relative to the fixation point. The yellow circles depict two example RFs locations. The red, green, and blue arrows indicate the radial-outward, radial-inward, and tangential directions with respect to the target location. **B.** The histograms illustrate the correlation between tuning curves for target alone vs target + one distractor calculated at 12 distractor positions, which were realigned based on the radial and tangential axes originating from the fixation point. Filled bars indicate statistically significant cases. In the polar plot, red and black data points compare the median values (red vertical lines) and proportion of significant cases (filled bars) from histograms of the matched directions, respectively.

In the “target + one distractor” condition, we studied responses to targets in the presence of a single distractor positioned at each of ten evenly spaced near surround locations on each trial (Figure 1C (iii) and see *Methods*). From these data for each neuron, we computed the correlation between target alone and target + one distractor at each of the ten locations. These correlation values underestimate the true correlation because they are based on a single repeat (25), but we can aggregate the data across neurons to look for population trends. We determined the radial/tangential axes based on the RF location of each neuron (Figure 4A), then realigned distractor locations onto these axes (see *Methods*) (Figure 4B).

Along the radial axis, the correlation in shape tuning between target alone and target + one distractor was significantly weaker for outward than inward position (compare 0° vs. 180° in the polar plot; median r: 0.29 vs. 0.56, Mann-Whitney U-test: p < 0.01; proportion of significant correlation cases: 18.0% vs. 29.2%), consistent with the anisotropy observed in psychophysical studies. Psychophysical studies have also demonstrated radial vs. tangential anisotropy (4, 10, 24), but we cannot address this point because we did not test the influence of a pair of distractors positioned radially vs. tangentially.

### Visual crowding in V4 cannot be explained by pooled encoding of nearby stimuli

One popular model to explain visual crowding posits that nearby stimuli within the receptive field of neurons in mid-level visual cortex may be pooled and encoded, e.g., in terms of their summary statistics (14–16). To test this hypothesis rigorously, we pursued two additional lines of inquiry. First, we asked whether enhancing the saliency of the target stimulus reverses crowding effects (Figure 5) reasoning that it *would not* if the effect was due to pooled encoding. Second, we asked whether metameric versions of crowded displays evoked similar responses (Figure 6) reasoning that it *would* if the representation was based on pooled image statistics.

**Figure 5.**
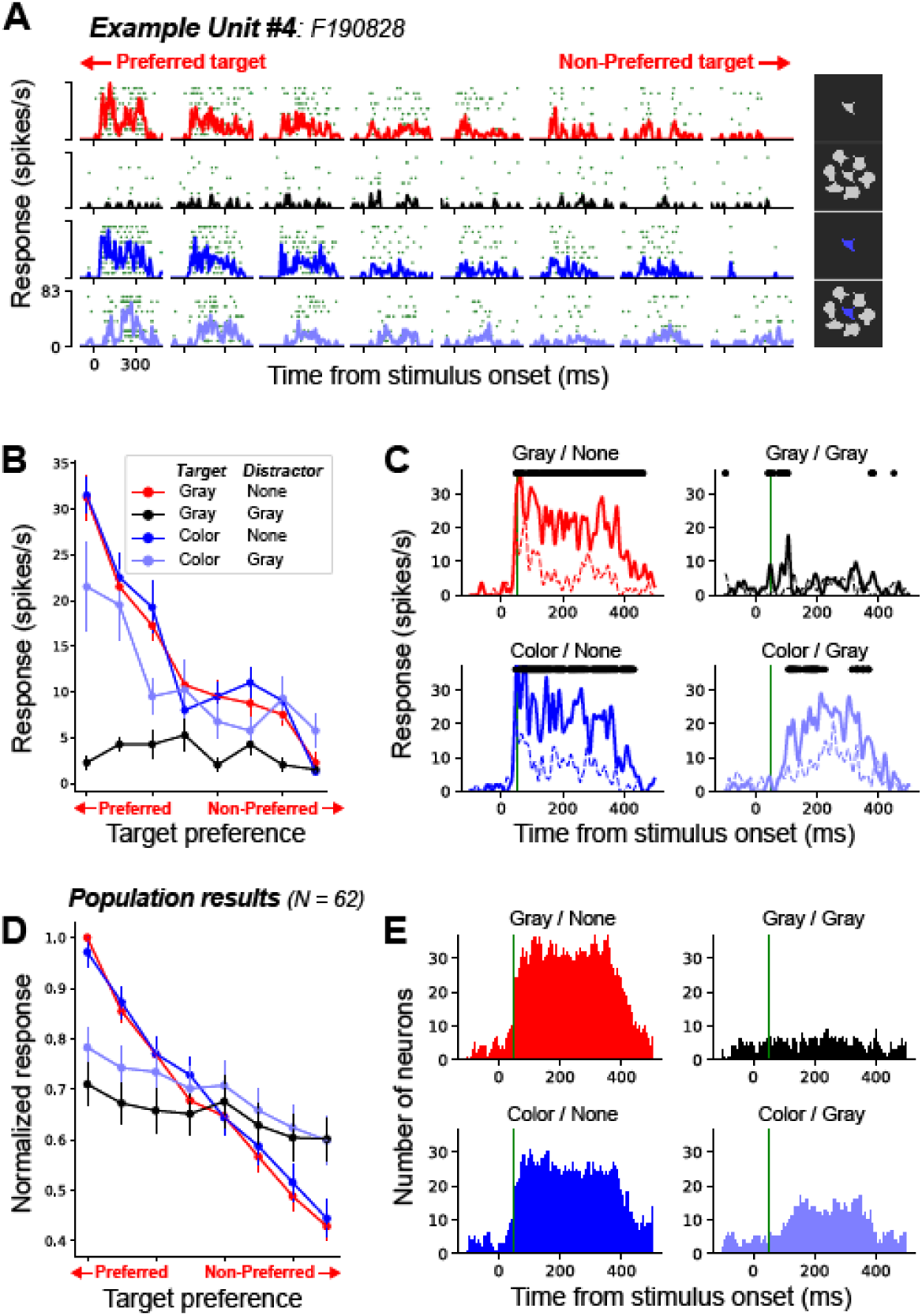
Effects of target saliency (color cue) on visual crowding. Target (gray or colored) appeared alone or in combination with six random distractors. **A**-**C.** Example unit response. **A.** raster plots with PSTHs. **B.** Target selectivity curves of the example unit for the four different conditions. **C.** Average PSTHs for the preferred (top 4) and non-preferred (bottom 4) targets. **D-E.** Population results. **D.** Average normalized tuning curves for target shape selectivity across conditions. **E.** The number of significantly modulated neurons as a function of time for each distractor condition. All conventions are as in Figure 2.

**Figure 6.**
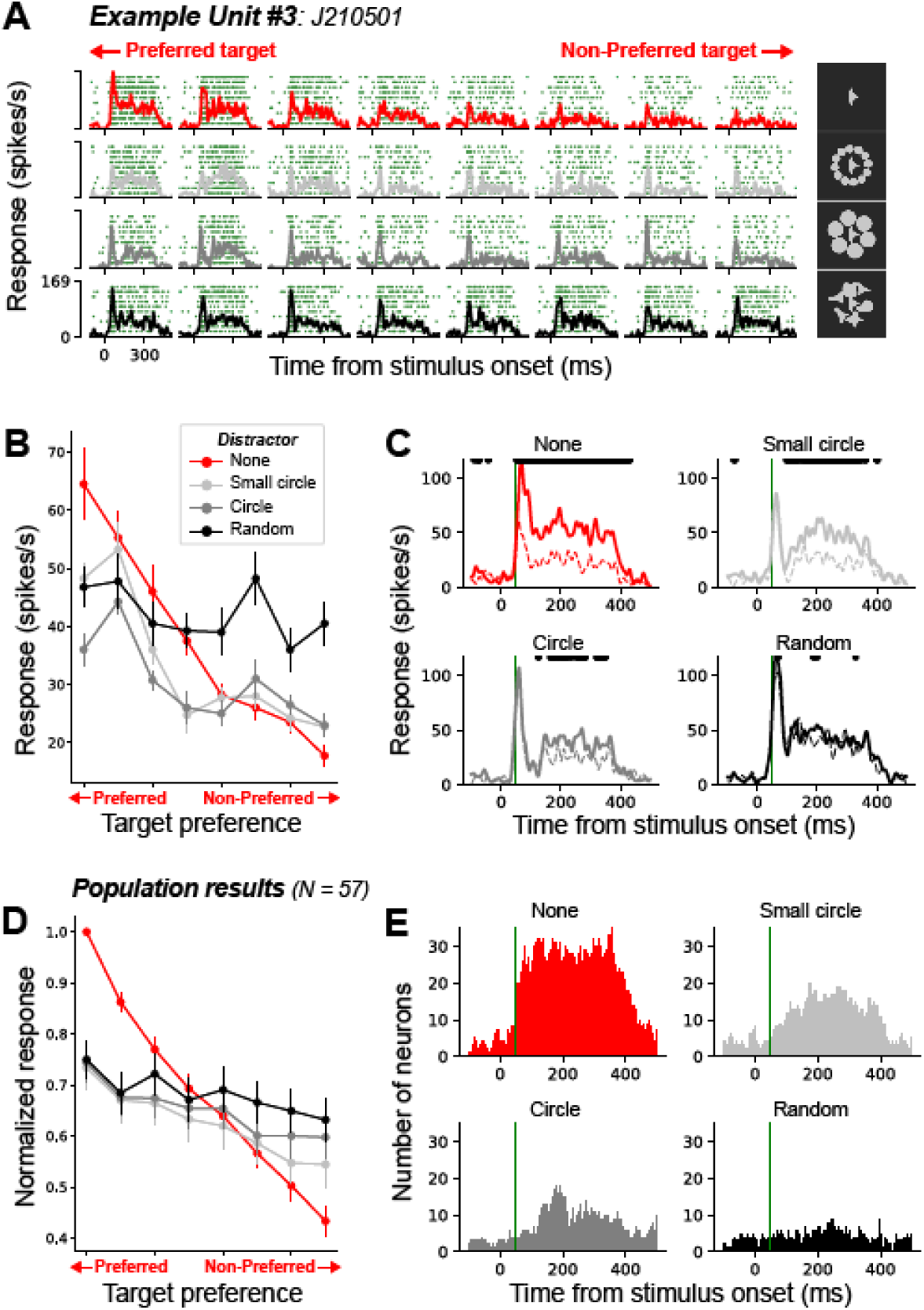
Effects of target saliency (shape, size cues) on visual crowding. Target appeared alone or in combination with 12 small circles, six circles, or six random distractors. **A**-**C.** Example unit response. **A.** Raster plots with PSTHs. **B.** Target selectivity curves of the example unit from four different conditions. **C.** Average PSTHs for the preferred (top 4) and non-preferred (bottom 4) targets. **D-E.** Population results. **D.** Average normalized tuning curves for target shape selectivity across conditions. **E.** The number of significantly modulated neurons as a function of time for each distractor condition. All conventions are as in Figure 2.

#### Enhancing saliency

In the preceding experiments, the distractors and target shape were achromatic, possessed bounding contours with varying curvature, and were sized to be ∼0.5 × RF diameter in linear extent (see Methods). Thus, the target stimuli did not stand out relative to the distractors. We used two strategies to enhance target saliency. First, the central target was defined by a chromatic contrast in addition to the luminance contrast, and we asked how this influenced the strength of responses and shape selectivity.

Figure 5 compares responses to achromatic (gray) and color target stimuli surrounded by 6 random shaped distractors. Importantly, gray and color targets were defined by the same luminance contrast, but the color targets were also defined by a chromatic contrast. For the example neuron in Figure 5A-C, the shape responses and tuning curves under the ‘target alone’ condition were very similar for the gray and color targets, indicating that shape selectivity was not affected by target stimulus color for this neuron. However, in the ‘target–distractor’ conditions, the results were markedly different. When the gray target was surrounded by gray distractors (black curves), target shape selectivity was completely abolished. In contrast, when the central target had a different color (light blue curves), the presence of surrounding distractors had little impact on shape selectivity (Figure 5B).

The population-averaged tuning curves of the 62 shape-selective neurons (62/135; 45.9%) reaffirmed that shape selectivity for color targets was more pronounced when compared to gray targets in the presence of multiple achromatic distractors (Figure 5D). While the effects were strong in some single units (as in Figure 5D), across the population, it was not strong as observed in the far distractor condition or the single distractor condition (light gray curves in Figure 2D and 3D). A more pronounced finding was that the onset of target shape selectivity was significantly delayed in the color target + gray distractor condition when compared to the color or gray target alone condition. This delay was consistently observed in both the example neuron data and the population data (Figure 5C and E).

In a second experiment, rather than modifying the target directly, we enhanced its saliency by presenting distractors of the same shape (all circles) that could be perceptually grouped together (26, 27). Similar to the randomly shaped distractors, these circle distractors were achromatic and positioned at a distance of 0.5 × RF diameter, but they were either the same size as the target (Figure 6A, 3^rd^ panel) or smaller (Figure 6A, 2^nd^ panel) further enhancing the saliency of the central target. The results obtained from an exemplar neuron (Figure 6A-C) demonstrate that shape selectivity is better preserved when all the distractors share a circular shape (correlation coefficient, r: 0.55 (random) vs. 0.79 (circle)), and this effect is further enhanced when the circles are smaller than the target shape (r = 0.91). Across the population (57/133; 42.9%), we observed an increase in the slope of the target shape tuning curve and enhanced shape selectivity as more salient cues were added to the target in cluttered scenes (shape cues → shape + size cues) (Figure 6D).

The analysis of the temporal dynamics of target shape selectivity once again highlighted notable distinctions between the target alone condition and the condition involving a salient target in a cluttered scene. For the example neuron (Figure 6C), the PSTHs across the different conditions showed a similar profile in terms of response onset and peak time, but striking contrast in terms of when significant differences emerged between preferred and nonpreferred responses. For target alone, significant differences emerged soon after response onset, but much later (after peak time) for salient targets presented in cluttered scenes (Figure 6C & E).

#### Comparison with metameric stimuli

To investigate the relationship between texture-like representation and visual crowding, we used a texture synthesis model to create synthetic images that differ physically from the originals (i.e., target + distractors) but have matched texture statistics within the RF (see *Methods*). If it is true that a single neuron in area V4 encodes a cluttered visual scene using texture-like summary statistics, then the neuronal responses to the originals will have a strong correlation with the neuronal responses to the synthetic images and relatively weak correlation with ‘target alone’ responses. Our results show that this prediction is not true. Responses to ‘target + distractors’ were better correlated with responses to ‘target alone’ than those to texture statistics matched synthetic (metameric) images (Figure 7A). Moreover, this finding was clearer when the target was salient (i.e., target + circle distractors) (Figure 7B). This indicates that V4 neurons encode the shape information of a salient target, which is not captured by the texture statistics model.

**Figure 7.**
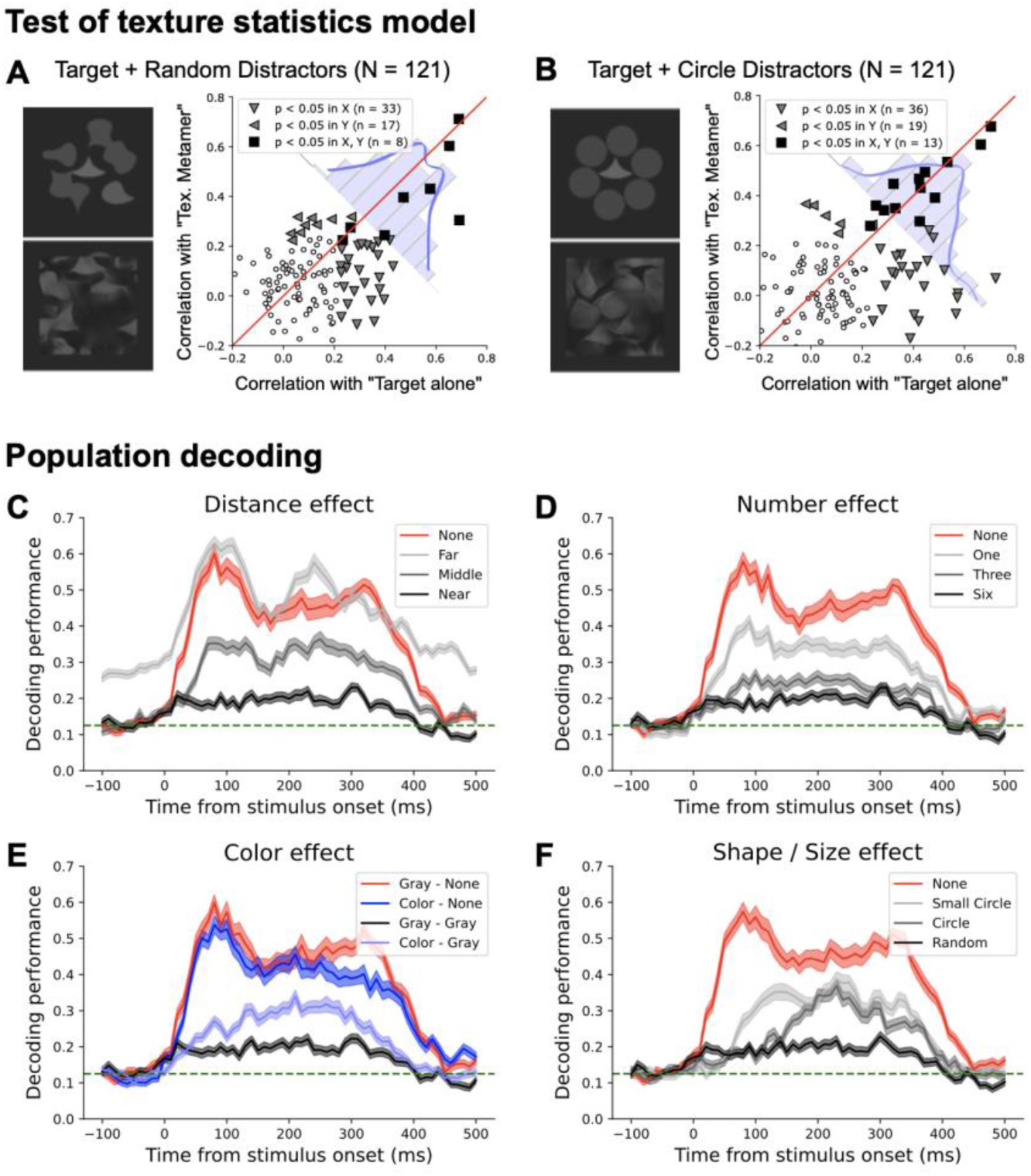
Test of texture statistics model and temporal dynamics of population decoding. **A-B.** Test of the texture statistics model for visual crowding. **A.** For each neuron, we computed two correlations: 1) the correlation between responses to ‘target + random distractor’ and ‘target alone’ stimuli, and 2) the correlation between responses to ‘target + random distractor’ stimuli and matched metamers. Population data are shifted below the diagonal suggesting that responses to target + random distractors are better correlated with the “target alone” condition (X-axis). Cells with a significant correlation (p < 0.05) in X-axis alone, Y-axis alone, or both are identified (see legend). **B.** The same analysis as in **A** for the ‘circle distractor’ condition in which the target is more salient. **C-F.** Population decoding. Population decoding performance plotted as a function time across different distractor conditions. In all four panels, target alone (red) and target + 6 near distractors (black) are identical. Decoding performance declines in the presence of distractors but time course varies across conditions. For the salient target conditions (light blue curve in **E**, gray curves in **F**) rise time and the maximum decoding performance time are delayed compared with target alone conditions (red and blue curves in **E**, **F**), but this is not the case for distance and number effects (gray curve in **C, D**). Green line indicates the chance level of target orientation decoding (0.125, 1 out of 8). Different colored lines represent different target-distractor configurations.

### Temporal dynamics of perceptual grouping

To begin to understand the time course of saliency related processes and the associated neural processes involved in reversing effects of crowding, we quantified the target shape decoding performance as a function of time using a 100 ms sliding window for the different classes of stimuli tested here. Since we studied all neurons with the same set of stimuli and every trial was unique (in terms of target and distractor placement combinations), we constructed a pseudo-population across all recording sessions. Using the far distractor condition as the training dataset, we first used PCA to derive a low-dimensional representation of the population data then used LDA to build a classifier for target stimulus orientation (see Methods). This classifier was then used to decode target orientation across all other stimulus conditions.

For the achromatic target alone condition, decoding performance (red, all panels) showed an immediate increase after stimulus onset, reaching its peak approximately 100 ms afterward. As expected, the decoding performance declined in the distractor distance (Figure 7C) and number (Figure 7D) conditions, but the dynamics were quite similar (compare gray and red lines, Figure 7CD). For the condition where we presented 6 distractors close to the target (black lines), decoding performance remained close to chance level.

However, under conditions where there are multiple distractors in close proximity to the target, the temporal dynamics of decoding performance varied markedly with saliency. When a target was salient by a color cue, decoding performance showed a delay in both the rise time and the time at which maximum performance was achieved, compared to the target alone conditions (Figure 7E). This was also the case when saliency was titrated by distractor shape/size cues (Figure 7F). This finding strongly supports the notion that the effects of target saliency and those of distractor distance/number are mediated by distinct neuronal mechanisms.

### Alternative model: saliency computation

Our findings strongly indicate the need for an improved model for visual scene encoding not only explains visual crowding effects but also incorporates the computation of visual saliency. Specifically, our results suggest that salient stimuli, defined as those stimuli with a high contrast in visual features relative to nearby stimuli, may be elevated and preferentially encoded in V4. Many researchers have proposed biologically plausible models for calculating visual saliency (28–30). Although these models may vary in specific details, they share common aspects: 1) early visual areas extract the most basic visual features of an image such as orientation, luminance, color, texture in parallel and pass the outputs to the next level area, 2) later stages integrate differentially weighted feature-based representations to create a saliency map. Here we incorporate this saliency map strategy to schematize an alternative model for processing of visual scenes with multiple shapes that could account for the observed crowding effects in area V4 (Figure 8). In the first processing stage, the visual input undergoes parallel processing through a group of low-level feature detectors (linear filtering), such as orientation, color, luminance, and texture. For our simulation, we utilized 64 conv1 layer filters from AlexNet (31), but more intricate models can be integrated, encompassing filters with different scales. The output of the linear filtering process (convolution of the input image and each of the 64 linear filters) produces the feature maps which are illustrated for the two example stimuli.

**Figure 8.**
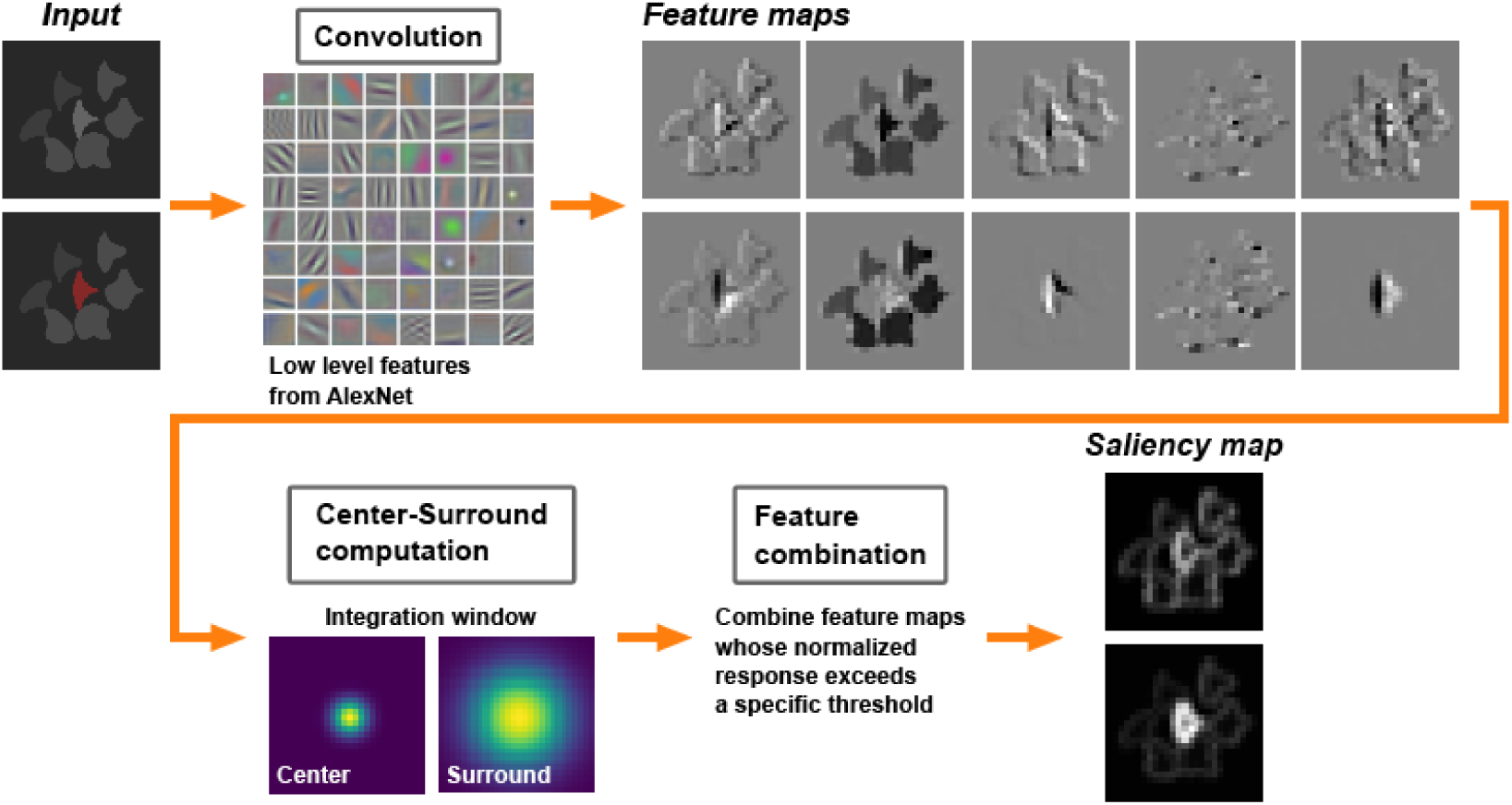
Hierarchical saliency computation model. Visual input is first processed in parallel by a set of low-level feature detectors (e.g., orientation, color, luminance, texture) in earlier visual areas. To focus on the central region of the visual scene, visual inputs (112 x 112 pixels) and feature maps (28 x 28 pixels) were cropped from the larger images of size 224 x 224 pixels and 55 x 55 pixels, respectively. Feature maps show the outputs from the first five filters from AlexNet. The next stage of processing performs a RF center-surround operation for each feature dimension and selectively combines only informative feature maps in which the RF center region is more strongly activated than its surround (see more details in the main text).

Next, consistent with prior work (28, 32, 33), we propose a within-feature surround normalization process which elevates high contrast features within each map. These normalized maps may then be thresholded, pooled, and form the input for shape selective computations in V4. Figure 8 shows our simulation results for two stimuli. In the case of a chromatic central target, feature maps based on chromatic contrast may elevate salience of the target stimulus compared to when the central target is also achromatic.

## Discussion

We investigated the neural correlates of visual crowding in macaque area V4. Our findings align with previous human psychophysics research, demonstrating that neuronal selectivity for object shape systematically declines with increasing distractor number, decreasing target–distractor distance, and exhibits spatial structure in its susceptibility to crowding. Importantly, our findings demonstrate that a salient target could mitigate the influence of nearby distractors, consistent with psychophysical results of uncrowding. To the best of our knowledge, our results provide the first comprehensive description of how various target–distractor configurations modulate V4 shape selectivity in single neurons. This delineates how simple scenes are encoded and its implications for both causing and alleviating crowding.

### Saliency trumps crowding

Traditionally, the predominant idea has been that adding distractors to a visual scene decreases the discriminability of a target (34–36). However, Herzog and colleagues recently demonstrated an “uncrowding effect” whereby crowding by flankers could be reversed by the appropriate placement of a larger number of flanking elements (37–39). Our own results are consistent with these findings. Even in highly cluttered visual scenes, the detrimental effects of distractors on V4 neuronal shape selectivity can be mitigated when target objects possess distinctive features, in terms of shape, size, or color, that set them apart from surrounding distractors. These results indicate that visual crowding effects observed in neuronal responses are consistent with psychophysical reports, in that they do not simply depend on target–distractor separation, but also on the spatial position of the distractors and the attributes of the distractors and target stimuli.

The uncrowding effect of Herzog and colleagues is thought to rely on different levels of perceptual grouping cues, ranging from low-level feature similarity between target and distractors to global context integration such as contour completion and may be closely linked to the computation of target saliency (39–41). However, the simple bottom-up saliency computation stream proposed in this study (Figure 8) does not incorporate global context information, which may be influenced by feedback signals from higher-order visual areas. Future research should investigate the role of feedback signals in perceptual grouping and the computation of target saliency.

The visual crowding effect, which relies on target saliency, shows similarities to the mechanisms that distinguish specific sounds from background noise. In both cases, the perceptual system encounters challenges when extracting and isolating relevant information in a cluttered or noisy environment. In the auditory domain, the detectability of a target sound in the presence of masking can be increased by adding sound energy that is frequency-distant from both the masker and the target (42, 43). This effect is observed when the remote sound and the masker share a consistent pattern of amplitude modulation, which is known as comodulation masking release (CMR). Interestingly, in our experiments, we observed a similar phenomenon where increasing the grouping cues among distractors enhances the saliency of the target and consequently reduces visual crowding.

### Encoding salient objects in V4

Many past studies have described in detail how single isolated stimuli— oriented bars, shapes and texture patches—modulate the responses of neurons in mid-level stages of the ventral visual pathway (44, 22, 45, 46). But how simple scenes composed of multiple objects are encoded and how that contributes to visual crowding is largely unknown.

One popular theory posits that neurons with peripheral receptive fields encode visual scenes in terms of texture-like summary statistics, integrating information across spatial regions that increase in size with eccentricity (15). Supporting this idea, a psychophysical study demonstrated that human observers struggle to discriminate images synthesized to have matched texture statistics at the receptive field sizes of area V2 in the ventral visual pathway (16). However, alternative perspectives argue that texture-based features alone are insufficient for representing the heterogeneous global structure of a scene (47, 48). In this study, we directly addressed the limitations of the texture summary statistics model in explaining the crowding effect. We found that the responses of V4 cells to a target surrounded by multiple distractors were more similar to the responses in the target alone condition, which had significantly different texture statistics, compared to the responses to metamers with matched texture statistics. This trend was particularly pronounced when the target was salient. Thus, our results provide further support to the hypothesis that V4 neurons encode segmented objects in a visual scene (21). When one of multiple objects is salient, that object is preferentially encoded across the V4 population. If multiple objects are equally salient, the individual objects may remain perceptually inaccessible due to limited processing capacity and the phenomenon of crowding ensues (26, 41).

### Visual processing stages that support (un)crowding

To uncover neural mechanisms underlying crowding, as in prior work (13, 49), we targeted area V4 for several critical reasons. RF sizes of V4 neurons are in good agreement with the sizes of crowding zones (e.g., 0.5 × Target eccentricity) (7, 50). V4 neurons encode diverse types of stimulus properties such as shape, texture, color, motion, disparity (21, 51). It is also well acknowledged that V4 responses are strongly modulated by both spatial and feature-based attention. V4 neurons exhibit flexible representation of visual stimuli depending on the feature relevant for the task being performed (52, 53). Therefore, it is optimal for computing target saliency. However, previous studies have shown that visual crowding effects are evident as early as V1 or V2 (54–56), suggesting that crowding may be based on activity at multiple levels throughout the visual processing hierarchy (4, 57, 58). Indeed, functional magnetic resonance imaging (fMRI) and electroencephalogram (EEG) investigations are consistent with this view. A recent study reported that surrounding distractors led to a greater impairment of neuronal discriminability for the orientation of a small target grating stimulus in V4 than V1 (13), and our preliminary data suggest that, unlike in V4, in anesthetized V2 recordings target saliency does not mitigate the detrimental effects of distractors on shape selectivity (59). Furthermore, the extent of visual crowding varies with perceptual experience (or training), and these experience-dependent changes have been reported to be highly specific to the training domain (60–62). This suggests that the phenomenon of visual crowding involves multiple brain regions specialized for distinct functions. Further studies with stimuli of different levels of complexity can improve our understanding of how different functions in different areas are impaired under the crowding phenomenon.

## Conclusion

In this study, we investigated the neural mechanisms of the visual crowding effect in macaque area V4, known for its significant role in object encoding. By manipulating diverse target-distractor configurations, we demonstrated that the visual system doesn’t rely solely on summary statistics to efficiently encode visual scenes with limited resources. Instead, it extracts salient features and processes them selectively. Future experiments are needed to gain deeper insight into how stimulus saliency is computed at various stages along the visual hierarchy.

## Materials and methods

### Animal preparation

Two healthy adult macaque monkeys (Monkey 1: male, 9.0 kg, 9 years old; Monkey 2: female, 6.0 kg, 9 years old) participated in the study. Animals had custom-built head posts anchored to the skull with orthopedic screws. A V4 recording chamber was placed over the left pre-lunate gyrus based on structural MRI scans. A craniotomy was performed in a subsequent surgery, 1-2 days before the first recording date. All animal procedures conformed to NIH guidelines and were approved by the Institutional Animal Care and Use Committee at the University of Washington.

During experiments, animals were seated in a primate chair in front of an LCD monitor (57cm away) and were required to hold their gaze within 1° of a small central fixation spot (0.1° diameter). As animals were engaged in this simple passive fixation task, a series of stimuli were presented in the visual periphery.

### Data collection

Recordings were performed using an epoxy-insulated tungsten microelectrode (PHC). The microelectrode was lowered into the cortex with a hydraulic microdrive (MO-97A; Narishige). Signals were amplified, bandpass filtered (150 Hz and 8 kHz) and digitized (sampling rate 32 kHz) with a Plexon MAP system (RASPUTIN v2 HLK3 System; Plexon Inc). Spike waveforms were sorted offline using principal component analysis (Offline Sorter; Plexon Inc). Time stamps of single unit spiking activity, eye positions (Eyelink 1000; SR research), and stimulus events (verified with photodiode signal) were stored at 1 kHz sampling rate for later analysis.

### Experimental procedures

Once a well-isolated single unit was identified, we first estimated the position of the receptive field (RF) using a hand-mapping procedure as animals were engaged in a passive fixation task (see *Animal preparation*). This was followed by an automated mapping procedure on a 7 x 7 grid (1 deg step) centered on the RF estimated by hand-mapping. For this experiment, we used a circular random dot motion patch (diameter: 1.4 deg) and presented the stimulus for 300 ms, 10 times at each of the 49 grid locations in random order. The revised RF center was estimated by fitting a 2D Gaussian to the measured responses (see *Receptive field shape estimation* below for more details) and the RF size was estimated based on the following equation (50) based on data from (22):

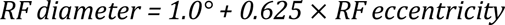

Following the RF mapping, the main experiment was performed to evaluate the effect of clutter on shape responses. While a monkey maintained fixation, a series of 3-4 stimuli (see *Clutter conditions* below) were presented, each for 300 ms with a 300 ms inter-stimulus interval, at the center of the RF. We studied the responses of 147 well-isolated V4 neurons with RF eccentricities within the central 10° of the visual field (Mean ± SD: 5.45 ± 1.80° for Monkey 1; 5.48 ± 2.05° for Monkey 2). For all neurons, we collected a minimum of 6 repetitions for each of the 111 stimulus conditions described below.

### Clutter conditions

To determine how surrounding clutter modulates shape responses and selectivity of V4 neurons, we measured the responses of each neuron to a central target shape in the presence of an array of surrounding distractors that were systematically varied across trials. Overall, we had a total of 111 stimulus conditions (see Figure 1A) where a central target shape was presented at one of eight rotations (0° – 315°, 45° steps) either alone or in combination with surrounding distractors that varied in number, distance from central target, and their saliency. The following paragraphs describe our target and distractor shape and clutter arrangements in detail.

#### Target and distractor shapes

The target and distractor shapes were chosen from a standard set of 2D stimuli constructed by a systematic combination of convex and concave boundary elements (Figure 1B). In previous studies (22, 45), this shape set has been used successfully to evoke a broad range of responses, facilitating the characterization of shape selectivity in V4 neurons tuned to different boundary features, e.g., convexities pointing up, concavities to the left, etc. For this study, we used one shape as the target stimulus (red box) across all recorded neurons. This shape has different convex and concave features along the boundary, and when presented at eight rotations (0° – 315°, 45° steps), could evoke a broad range of responses from individual neurons (see *Results*). The target shape (achromatic; luminance: 20 cd/m^2^) was presented at the center of the RF and its size was adjusted to 0.5 × RF diameter. The luminance of the achromatic background was set to 2.4 cd/m^2^. The distractor shapes were randomly chosen from the same shape stimulus set. They were achromatic and their luminance was randomly set to one of 5.7, 9.8, or 15.4 cd/m^2^. Distractors were arranged around the target stimulus at varying distances and sizes as noted below.

#### Target-distractor distance

Target shapes were surrounded by distractors positioned at three radial distances from the target: near, middle, or far (Figure 1C (ii)). In the near distance condition, six distractors were evenly spaced and positioned at a radial distance equal to the RF radius of the neuron under study. For middle and far distance conditions, the center-to-center distances between the target and distractors were set to 2 × and 3 × radius of RF, respectively. To keep the distance between distractors (i.e., density of distractors) constant across the three distance conditions, we used 12 and 18 distractors for the middle and far distance conditions, respectively. Distractor shapes were randomly chosen for each repetition, but they were the same across days. Thus, every neuron in our dataset was subjected to the same array of target + distractor stimuli that were re-positioned and scaled to match RF center and size. To avoid stimulating the exact same position with the distractor stimuli, distractors were evenly spaced around the circle, but the precise angular positions were randomly chosen. In addition, we introduced a minor spatial jitter in both the horizontal and vertical positions of individual distractor positions. This jitter amounts to approximately 10% of the size of the distractor stimulus.

#### Number of distractors

To determine how the number of distractors influences responses, we presented the target shape surrounded by one, three, or six distractors (Figure 1C (iii)). The distance between the central target and surrounding distractors was fixed as the radius of the RF. Distractors were evenly spaced along the angular dimension (e.g., 120° apart for three distractors, 60° for six distractors) but their angular positions and shapes were randomly chosen for each trial with a small spatial jitter as described above. The angular positions were carefully calibrated across ten trial iterations to ensure uniform stimulation of the peripheral region of the RF. All neurons were tested with the same target-distractor arrangements.

#### Target saliency – color, shape, size

To determine whether target saliency, i.e., target contrast relative to distractors, resists effects of crowding, we compared responses to the target + distractor arrays in the near distance condition discussed above with three conditions where target saliency was enhanced using different cues: *color*, *shape*, and *size*. In the ***color*** saliency set (Figure 1C (iv)), the distractors were the same as in the near condition, but the target stimulus was defined by a chromatic contrast as well; its luminance contrast was the same as that of the achromatic target (20 cd/m^2^) in the near condition. In the ***shape*** saliency set (Figure 1C (v), 2^nd^ panel), all distractors were circles to facilitate distractor grouping. In the ***shape* + *size*** saliency set (Figure 1C (v), 1^st^ panel), the distractors were small circles: 12 small circles that were half the size of the target (0.5 × target size).

### Visual crowding models

#### Metameric stimuli: Texture statistics model

One prominent hypothesis posits that crowding results from encoding the visual image patch within the RF in mid-level visual cortex in terms of texture-like summary statistics (14–16). To rigorously test whether neuronal responses to the clutter displays in our main experiment can be characterized as encoding pooled texture statistics, we created metameric stimulus images for a subset of our stimuli using the Portilla–Simoncelli texture statistics model (63) (see examples in Figure 1C (vi)). The model extracts 740 texture synthesis parameters from each stimulus image, then iteratively modifies a white-noise image until its parameters are matched with those from the original stimulus image. We directly compared responses to the original source with those to the synthesized images that share pooled texture statistics. We performed this comparison for three of the stimulus conditions where we presented 6 random shape distractors, 6 circle distractors or 12 small circle distractors all at a distance of RF radius from the target stimulus (see Figure 1A).

#### Saliency computation model

We built a three-stage model to simulate how preferential encoding of salient stimuli might arise in mid-level cortical stages. The visual input for the model were the stimulus conditions discussed above, rescaled to 224 × 224 pixels. The first processing stage of the model extracts basic visual features using the first convolutional layer (Conv 1) of AlexNet, a deep neural network which was pre-trained to classify images into 1000 object categories (31). This stage convolves the input image with 64 filters of Conv1, and the resulting feature maps are represented with dimensions of 55 × 55 pixels each. The critical next step is to combine multiple feature maps into a single saliency map. Drawing from prior work (28, 32, 33), we quantified saliency as the absolute difference in intensity between the center and the surrounding regions. The center region and surround region were defined using spatially overlapping 2D Gaussian functions, with a three-fold difference in spatial scale (σ_center_ = 2 pixels, σ_surround_ = 6 pixels). Then, only those maps with high center-surround difference exceeding a threshold (activation from the center region > 2 × activation from the surround region) were linearly combined.

### Data analysis

#### Time course of spiking responses

On each trial, we extracted spike trains (1 ms bin) aligned on stimulus onset. Peri-stimulus time histograms (PSTHs) were constructed by averaging spike trains across multiple repetitions and convolving with a Gaussian kernel (σ = 5 ms).

#### Average responses of single neurons

For each stimulus condition, we quantified the average response magnitude by counting spikes within a window from 0 to 400 ms after each stimulus onset (to include both onset and offset responses of V4 neurons) and averaging across multiple repetitions.

#### Quantification of distractor modulation

To quantify how the distractors that surround the central target modulate responses to the target stimulus, we computed a distractor modulation index, given by:

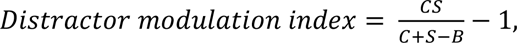

where 𝐶𝑆, 𝐶, 𝑆, 𝐵 represent neuronal responses to target–distractor stimulus configuration, center target stimulus alone, surround distractors alone, and baseline (i.e., no stimulus) conditions, respectively. The denominator indicates the response level estimated by linear summation of center target alone and surrounding distractors alone conditions. Negative values indicate suppressive modulation.

#### Modulation of shape selectivity by distractors

Shape selectivity of individual neurons was assessed by performing a one-way analysis of variance (ANOVA) of the responses to the eight rotations of the target shape presented in isolation. Neurons that showed significant (p < 0.05) shape selectivity (73/147; 49.3%) based on this one-way ANOVA were subjected to the following additional analyses. To quantify the effects of distractors on neurons’ shape discriminability, we evaluated the similarity of shape tuning between target alone condition and each of target–distractor conditions by calculating a Pearson’s correlation coefficient between them. A significant correlation coefficient implies that the shape selectivity survived despite the presence of distractors.

#### Average shape tuning curves across the population

To construct average shape tuning curves across the population for each clutter condition, we first normalized the tuning curves from each neuron by the responses to the most preferred target shape from the ‘target alone’ condition and averaged the normalized tuning curves across all neurons.

#### Temporal dynamics of target shape selectivity

To determine the timing of target shape selectivity in each of the distractor conditions, we divided responses into two groups (4 preferred targets vs. 4 non-preferred targets) based on target preference determined from target alone condition and asked when responses between the two groups significantly deviated from each other using the Mann–Whitney U test (p < 0.05) within a 30 ms sliding window (moving in 1 ms steps).

#### Receptive field shape estimation

We sought to examine whether V4 neuron responses exhibit anisotropies in the effect of clutter as a function of the distractor position relative to the RF center. For each neuron, we obtained the RF map from spiking responses elicited during the automated RF mapping paradigm (see *Experimental procedures*). We then fitted a two-dimensional elliptical Gaussian function to the map to determine the center, size (full width at half maximum (FWHM) = 2.355 × 𝛔), and the angle of the elongated axis of the RF.

A two-dimensional elliptical Gaussian function is expressed as

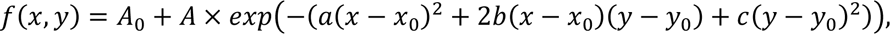

where 𝐴_0_ is a constant, 𝐴 is the scale factor, 𝑥_0_ and 𝑦_0_ are the center of the Gaussian along 𝑥 and 𝑦 axis, respectively.

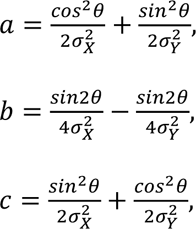

parameters 𝑎, 𝑏, 𝑐 are set as above, where 𝜃 represents the counter-clockwise rotation angle of the Gaussian blob. A value of 0 for 𝜃 indicates that the blob is pointing to the right.

#### Realigning distractor positions based on radial and tangential axes from the fixation point

To explore whether V4 neurons represent the anisotropy observed in visual crowding, we determined the radial and tangential axes for each neuron based on their RF locations. The radial axis is defined by the line connecting the RF center to the fixation point and the tangential axis is orthogonal to it. Subsequently, we realigned the 10 single distractors’ positions onto these established axes. The original positions of the distractors were spaced 36 degrees apart, but to accurately reflect the orthogonal radial and tangential axes, we divided the repositioned positions into 12 overlapping bins, each spanning 60 degrees and spaced 30 degrees apart.

#### Population decoding analysis

To analyze how target shape processing in V4 neurons evolves over time under different target–distractor conditions, we computed the population decoding performance as a function of time using a 100 ms sliding window. Using the shape selective neurons identified above (see *Modulation of shape selectivity by distractors*), the analysis procedure was as follows. We used principal component analysis (PCA) on the single trial responses to target + far distance distractor condition to derive a low-dimensional representation of the population response (five principal components (PC), explained variance = 35 %).

Single trial responses from all other target–distractor conditions were projected onto the space defined by these five PCs. A linear discrimination analysis (LDA) model was fitted using the low-dimensional representation of target + far distance distractor condition as the training data. The fitted LDA model was used to predict the target shape across all the other target–distractor conditions (chance level = 0.125). This procedure was repeated 100 times, involving the random selection of 30 neurons without replacement in each iteration.

### Experimental design and statistical analysis

Details of experimental procedure and visual stimuli are described above (see *Data collection, Clutter conditions*). The shape selectivity of individual neurons for the eight target shapes was assessed by conducting a one-way ANOVA. The strength of the linear relationship between pairs of variables (e.g., shape selective responses from ‘target alone’ and ‘target + distractor’ conditions) was assessed by Pearson’s correlation coefficient.

Wilcoxon signed-rank test was used for paired comparison of correlation coefficients or modulation indices between clutter conditions in each neuron. Independent group comparisons were performed using a non-parametric Mann–Whitney U test. A p-value of 0.05 or less was considered significant.

### Data and software availability

The data and analysis code that support the findings of this study are available from the corresponding author upon request.

## Author contributions

T.K. and A.P. contributed to conception and design of the experiments. T.K. conducted data collection. T.K. analyzed the data. T.K. and A.P. wrote the manuscript. T.K. and A.P. approved the final version.

## Acknowledgement

The authors are grateful to Rohit Kamath, Dr. Anjani Chakrala, and Dr. Dina Popovkina for providing helpful discussions and comments on the manuscript, and Amber Fyall for assistance with animal training. This work was supported by NEI Grant R01 EY018839 to A.P.; NEI Center Core Grant for Vision Research P30 EY01730 to the UW; NIH/ORIP Grant P51 OD010425 to the WaNPRC.

